# The epidemiological signature of pathogen populations that vary in the relationship between free-living survival and virulence

**DOI:** 10.1101/2020.06.08.141366

**Authors:** Lourdes M. Gomez, Victor A Meszaros, Wendy C. Turner, C. Brandon Ogbunugafor

## Abstract

The relationship between parasite virulence and transmission is a pillar of evolutionary theory that has specific implications for public health. Part of this canon involves the idea that virulence and free-living survival (a key component of transmission) may have different relationships in different host-parasite systems. Most examinations of the evolution of virulence-transmission relationships—theoretical or empirical in nature—tend to focus on the evolution of virulence, with transmission a secondary consideration. And even within transmission studies, the focus on free-living survival is a smaller subset, though recent studies have examined its importance in the ecology of infectious diseases. Few studies have examined the epidemic-scale consequences of variation in survival across different virulence-survival relationships. In this study, we utilize a mathematical model motivated by aspects of SARS-CoV-2 natural history to investigate how evolutionary changes in survival may influence several aspects of disease dynamics at the epidemiological scale. Across virulence-survival relationships (where these traits are positively or negatively correlated), we found that small changes (5% above and below the nominal value) in survival can have a meaningful effect on certain outbreak features, including the *R*_*0*_, and the size of the infectious peak in the population. These results highlight the importance of properly understanding the mechanistic relationship between virulence and parasite survival, as evolution of increased survival across different relationships with virulence will have considerably different epidemiological signatures.

## INTRODUCTION

The interactions between the life history of a pathogen and the environment in which it is embedded drive the evolution of virulence. These interactions thus dictate both the experience of disease at the individual host level, and the shape of disease dynamics in host populations [1,2]. The nature of the interaction between virulence and transmission has been the object of both theoretical and empirical examination [2-8]. Free-living survival, here defined as the ability of a pathogen to persist outside of its host, is one of many transmission life history traits associated with virulence. The relationship between the two varies between host-pathogen types and different environments [4,8-10].

Several hypotheses serve as canon in the evolution of virulence, theorizing on its relationship to transmission traits. The *Curse of the Pharaoh* hypothesis--named after a tale about a mythical curse that torments individuals who dig up tombs of Egyptian pharaohs [11] suggests that if a parasite has high free-living survival, then it is far less dependent on its host for transmission, and consequently, will have no evolutionary incentive to decrease virulence [2,4,12]. Spreading through a host population with high mortality can be supported by persisting in the environment until the arrival of new hosts or the recovery of the host population from survivors. That is, any presumptive selection on beneficence would be relaxed: parasites can detrimentally affect the health of hosts at no cost to transmission, because most of their life cycle is spent outside of a host. Previous studies support a positive correlation between free-living survival and mortality per infection (a common proxy for virulence) [13].

Alternatively, the “tradeoff” hypothesis suggests that there is some intermediate level of parasite virulence [3,6,14] that is optimal for a given setting. In this scenario, too high of virulence kills host and parasite, and too low of virulence leads to a failure to transmit. Applying this hypothesis specifically to free-living survival would suggest that selection for increased free-living survival should come at the expense of virulence (producing a pathogen that is less harmful to the host). Mechanistically, as a consequence of increased adaptation to a non-host environment, a virus may be less fit to replicate inside a host [9,15]. For example, a more robust viral capsid may help to survive harsh environmental conditions, but may make it more difficult to package RNA/DNA [15]. More generally, the tradeoff hypothesis can be framed in the context of a life history tradeoff: investment in certain parts of the life cycle often comes at the expense of others [2,16].

Theoretical studies have explored varying evolutionary relationships between heightened virulence and extreme pathogen longevity [4, 5, 12, 17-19]. One critical component of these studies revolves around whether or not virulence evolves independently of free-living survival. For example, some models have argued [4] that pathogen virulence is independent of its survival under a set of conditions: when the host-pathogen system is at an equilibrium (evolutionary and ecological), if host density fluctuates around an equilibrium, or if turnover of the infected host population is fast relative to the pathogen in the environment. However, if the host-pathogen system is at disequilibrium and if the dynamics of propagules in the environment are fast compared to the dynamics of infected hosts, then virulence is, as hypothesized, an increasing function of propagule survival [4]. Kamo & Boots [17] examined this hypothesis incorporating spatial structure in the environment using a cellular automata model, and found that if virulence evolution is independent of transmission, then long-lived infective stages select for higher virulence. However, if there is a trade-off between virulence and transmission, there is no evidence for the Curse of the Pharaoh hypothesis, and in fact, higher virulence may be selected for by shorter rather than long-lived infectious stages. Further, the evolution of high virulence does not have to occur solely through a transmission-virulence tradeoff. Day [18] demonstrates how pathogens can evolve high virulence, and even select for traits to kill the host (e.g., toxins), if pathogen transmission and reproductive success are decoupled. These studies emphasize the context-dependence of virulence-survival relationships. Understanding where in the relationship between virulence and survival a given pathogen population lives in a specific setting may allow one to understand how virus evolution will manifest at the level of the epidemic.

In this study, we examine the epidemic consequences of different virulence-survival relationships—direct vs. inverse proportionality— in a viral disease with an environmental- transmission component. In order to measure how pathogen survival influences disease dynamics, we included an “environmental compartment” in our model, which represents “contaminated” environments that act as a reservoir for persisting pathogens, causing disease spread when they come in contact with susceptible individuals (infection via “fomites”) [20, 21].

We find that the identity of the virulence-free-living survival relationship (e.g. positive vs. negative) has distinct implications for how an epidemic will unfold. Some, but not all, features of an outbreak are dramatically influenced by the nature of the underlying virulence-survival relationship. This indicates that signatures for evolution (adaptive or other) in a pathogen population will manifest more conspicuously in certain features of an outbreak. We reflect on these findings in light of their theoretical implications on the evolution and ecology of infectious disease, and for their potential utility in public health interventions.

## METHODS

### Model motivation and application

The mathematical model explored in this study is adapted from a recent one developed to investigate environmental transmission of SARS-CoV-2 during the early-stage outbreak dynamics of COVID-19, with parameter values based on fits to actual country outbreak data [26]. In this study, we utilize this model to examine questions about the evolution of free-living survival. While the phenomenon we examine is a very relevant one that manifests in the actual, biological world, we want to emphasize that none of the methods or results in this study are intended to apply to the active COVID-19 pandemic. This study is an attempt at responsible theoretical biology, with data-informed models and inferences that are germane to the natural world. However, we do not support the extrapolation of these findings to any particular aspect of COVID-19, nor should they inform a policy or intervention.

### Model Description

The model is implemented via a set of ordinary differential equations, defined by Equations 1.1 − 1.6. It implements viral free-living survival via the “Waterborne Abiotic or other Indirect Transmission (WAIT)” modelling framework, coupling individuals and the pathogen within the environment [27, 28].

Within the model, a β_w_ term allows for individuals to become infected via viral pathogen deposited in the environment, and terms σ_A_ and σ_I_ allow asymptomatic and symptomatic individuals to deposit pathogen into the environment, respectively. Adapted from the more traditional SEIR (susceptible-exposed-infected-recovered), the SEAIR-W (susceptible-exposed-asymptomatic-infected-recovered-WAIT) model interrogates the consequences of the two hypotheses outlined above, while representing the dynamics of a very relevant disease system (SARS-CoV-2) that includes an asymptomatic infectious population. While asymptomatic transmission remains a somewhat contentious topic, several studies have suggested that it plays a role in the spread of disease [22-24]. Though environmental transmission of SARS-CoV- 2 remains a controversial topic, it is plausible that asymptomatic individuals may spread disease through frequent contact with the environment, thus increasing the proportion of virus that is free-living [25].

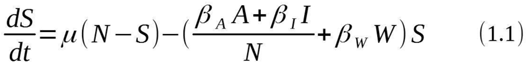

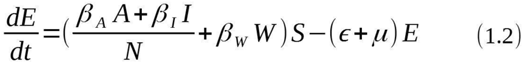

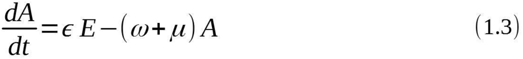

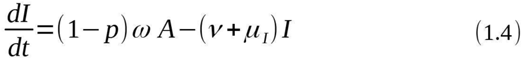

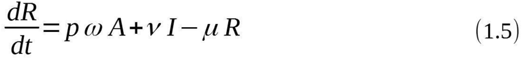

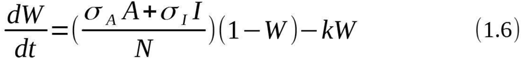

Figure 1 depicts the compartmental diagram for the model. The direction of the arrows corresponds to the flow of the individuals and the pathogen through the system. Note that individuals can move from the asymptomatically infected compartment to the recovered compartment, what we call a “mild track”. The dashed arrows represent the WAIT coupling to the environment. The model is inspired by a model developed to interrogate environmental transmission of SARS-CoV-2 [26].

**Figure 1.**
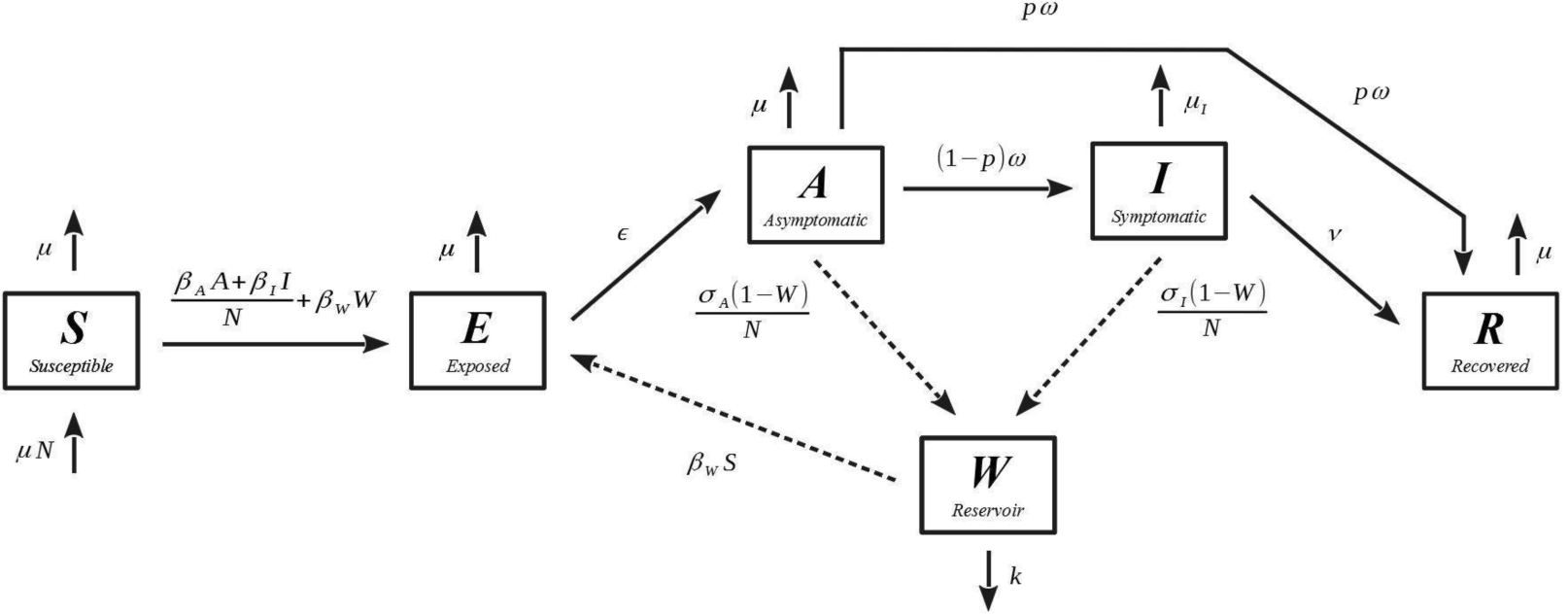
Compartmental diagram of the SEAIR-W model: Here we present the compartmental diagram for the model system highlighted in this study. The model is similar to a mathematical model used to interrogate environmental transmission of SARS-CoV-2 (see [26]).

### Simulations of outbreaks

The system was numerically integrated using the “odeint” solver in the Scipy 1.4 - the Python scientific computation suite [29]. The simulations track the populations for each of the compartments listed in Figure 1. Each model run occurred over 250 days, which amounts to over 8 months of the epidemic or 5x the peak of the outbreak. This length of time is consistent with the antecedent SARS-CoV-2 model [26] long enough for the dynamics of the system to manifest. Note, however, that for this study, we are especially interested in the early window of an outbreak, the first 30 days. We focus on this window because this is the window that best captures the underlying physics of an epidemic, as 30 days is often before populations have been able to adjust their individual behaviors. The code constructed for the analysis in this study is publicly available on github: https://github.com/OgPlexus/Pharaohlocks

### Population definitions & parameter values

Table 1 outlines the definitions of each population, and provides the initial population values used for all simulations conducted in this study. The nominal parameter values used are defined in Table 2. The initial values are drawn from the aforementioned COVID-19 outbreak study, derived from empirical findings and country-level outbreak data [26].

**Table 1.**
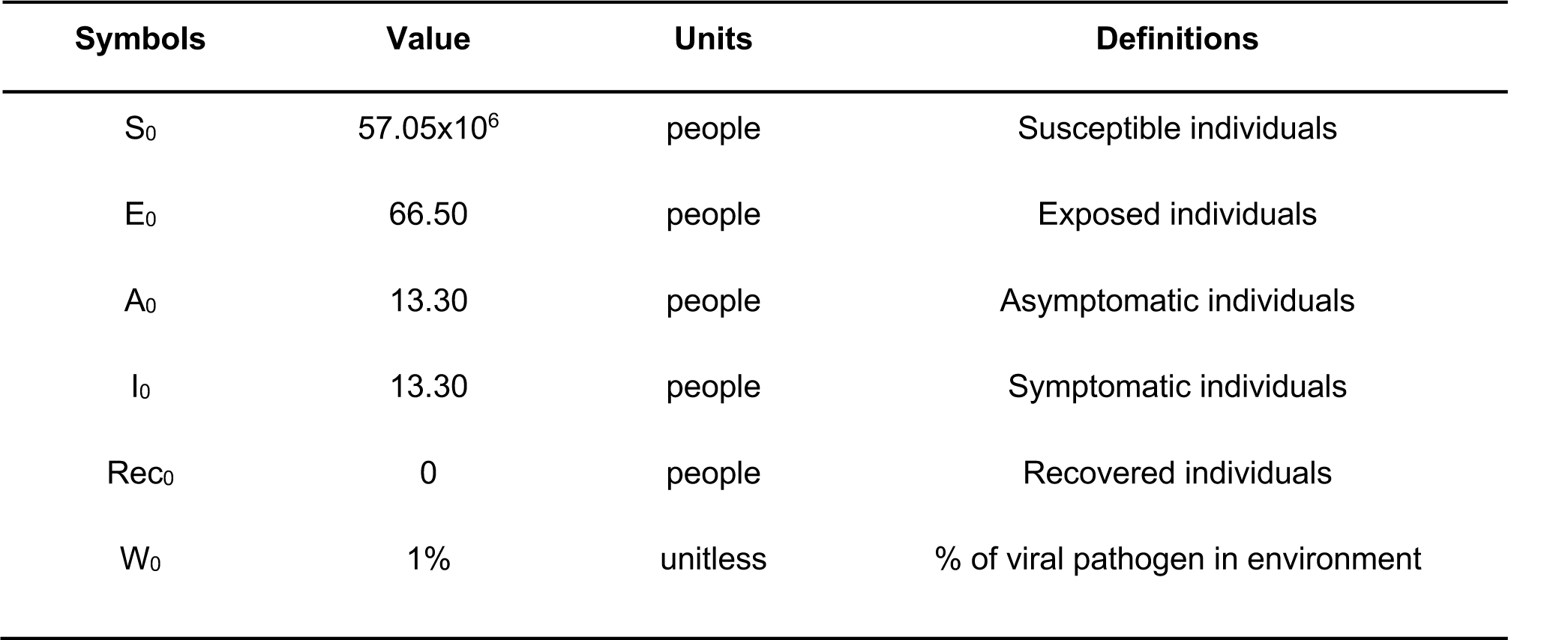
Definitions and initial values for the populations represented by each compartment. The values here are the averages of the model values across all the countries developed in a prior COVID-19 study [26].

**Table 2.** Definitions for the nominal parameter values used in this study. Parameter values were developed from empirical findings and country-level data, as discussed in another study [26].

| Symbols | Values | Units | Definitions |
| --- | --- | --- | --- |
| $\mu$ | 1/(80.3 * 365) | 1/day | Natural death rate (Reciprocal of the upper bound of average human lifespan) |
| $\mu_i$ | 0.00159 | 1/day | Infected death rate<br>(Natural death rate + disease induced death rate) |
| $\eta = (\omega - \varepsilon^{-1})$ | 5.5 | days | SARS-CoV-2 incubation period |
| $1/\omega$ | $\eta - \varepsilon^{-1}$ | days | Rate of transfer from asymptomatic to symptomatic |
| $\nu$ | 0.0305 | 1/day | Recovery rate (Average of 3 to 6 weeks) |
| $p$ | 95.6% | percent | Percent that move along the “mild” recovery track |
| $k$ | 0.649 | 1/day | Waning virus rate in environment (using average of all material values, wood, steel, cardboard, plastic) |
| $\beta_a$ | 0.550 | 1/day | (Contact rate of people with people) x (transmission probability of people to people by A-person) |
| $\beta_i$ | 0.491 | 1/day | (Contact rate of people with people) x (transmission probability of people to people by I-person) |
| $\beta_w$ | 0.031 | 1/day | (Contact rate of person with environment) x (transmission probability of environment to people) |
| $\sigma_a$ | 3.404 | 1/day | (Contact rate of person with environment) x (probability of shedding by A-people to environment) |
| $\sigma_i$ | 13.492 | 1/day | (Contact rate of person with environment) x (probability of shedding by I-people to environment) |
| $1/\varepsilon$ | 2.478 | days | Average number of days before infectiousness |

### Virulence definition

In this study, we define virulence as a set of parameters that uniformly increase the rate or probability of causing symptomatic disease, or the severity of those symptoms (including death). Our definition is more comprehensive than many other models of parasite virulence (e.g., [4,13]), which tend to focus on a single aspect of the natural history of disease associated with harm to a host (e.g., the fitness consequences of an infection on the host population, or the case fatality rate). Instead of having to justify a definition built around a single term (e.g. the term associated with fatality), we took a collective approach to defining virulence through all terms that foment the viral-induced onset of symptomatic disease, and death. This more flexible definition allows for the reality of pleiotropic effects in viral pathogens, where adaptations can have multiple effects on the natural history of disease [2,30]. Our definition of virulence emphasizes terms that influence host wellness and/or are symptoms of disease. This iteration of virulence used in the study also undermines the potential for overly-weighting only one or a small number of parameters under a large umbrella of virulence. Because so many varying definitions exist for virulence, we have also performed calculations according to a different definition of virulence, one that exclusively considers terms that have a detrimental direct effect on the host, and neither of the terms that reflect symptoms of severe disease (*σ*_*a*_ *and σ*_*I*_). These calculations can be found in the Supplemental Material.

The collection of parameters that we use to define virulence are as follows: the infected population death rate (*μ*_I_), the incubation period of SARS-CoV-2 (*η*), the rate of transfer from asymptomatic to symptomatic (1/ω), the infected population recovery rate (v), the percent of individuals that move from the asymptomatic to the recovered compartment without showing symptoms (the “mild” recovery track, p), the contact rate of people with people ***X*** transmission probability of people to people by an asymptomatic individual (β_A_), the contact rate of people with people ***X*** transmission probability of people to people by asymptomatically infected person (β_I_), the contact rate of people with environment ***X*** probability of shedding by asymptomatic individual to the environmental (*σ*_A_), the contact rate of people with the environment ***X*** probability of symptomatically infected individuals shedding into the environment (*σ*_I_), and the average number of days before infection (1/ ε).

Table 3 outlines the direction in which each of the virulence-associated parameters are modulated as virulence decreases or increases. An up arrow (↑) indicates the parameter increases (by an equivalent percent) when the percent virulence is changed. A down arrow (↓) indicates the parameter decreases (by an equivalent percent), when the percent change in virulence is applied. Changes in virulence are then defined, in this study, as an equivalent uniform (percent) change in each of the parameters listed above. For the purposes of our study, we modify virulence by changing all parameters associated with virulence by 5%. It would also be easy to disambiguate virulence into changes in individual subcomponents, however, that is not the focus on the current study.

**Table 3.** Virulence parameters. This is a list of uniformly modulated parameters, and the direction in which they change, when virulence is increased or decreased. When virulence changes, an up arrow (↑) indicates the parameter is increased (by an equivalent percent), and a down arrow (↓) indicates the parameter is decreased (by an equivalent percent).

| Symbols | Definition | Virulence Increased | Virulence Decreased |
| --- | --- | --- | --- |
| $\mu_i$ | Infected death rate<br>(Natural death rate + disease induced death rate) | ↑ | ↓ |
| $\eta = (\omega - \varepsilon^{-1})$ | SARS-CoV-2 incubation period | ↓ | ↑ |
| $1/\omega$ | Rate of transfer from asymptomatic to symptomatic | ↑ | ↓ |
| $\nu$ | Recovery rate (Average of 3 to 6 weeks) | ↓ | ↑ |
| $p$ | Percent that move along the “mild” recovery track | ↓ | ↑ |
| $\beta_A$ | (Contact rate of people with people) x (transmission probability of people to people by A-person) | ↑ | ↓ |
| $\beta_I$ | (Contact rate of people with people) x (transmission probability of people to people by I-person) | ↑ | ↓ |
| $\sigma_A$ | (Contact rate of person with environment) x (probability of shedding by A-people to environment) | ↑ | ↓ |
| $\sigma_I$ | (Contact rate of person with environment) x (probability of shedding by I-people to environment) | ↑ | ↓ |
| $1/\varepsilon$ | Average number of days before infectious | ↓ | ↑ |

### Survival definition

Survival is defined as the set of parameters that when uniformly modulated, increase the pathogen’s probability of surviving the outside environment and successfully infecting a new host [2]. In our model this includes both the waning virus rate in the environment (k) and the contact rate of an individual with environment ***X*** transmission probability of environment to people (β_w_). Table 4 outlines the direction (increasing or decreasing) in which these parameters are modulated when survival is decreased or increased. Within both models a (percent) change in survival is defined as an equivalent uniform (percent) change in the survival parameters.

**Table 4.** Survival parameters. This is a list of parameters that are uniformly modulated, and the direction in which they change, when survival is increased or decreased. An up arrow (↑) indicates the parameter is increased by some percent, when the equivalent (percent) change in survival is applied. A down arrow (↓) indicates the parameter is decreased by some percent, when the equivalent (percent) change in survival is applied.

| Symbols | Definition | Survival Increased | Survival Decreased |
| --- | --- | --- | --- |
| $k$ | Waning virus rate in environment (using average of all material values, wood, steal, cardboard, plastic) | ↓ | ↑ |
| $\beta_w$ | (Contact rate of person with environment) x (transmission probability of environment to people) | ↑ | ↓ |

Throughout this study, the impact of changes in virulence and survival (and the relationship between these traits) are assessed with respect to the following four epidemic metrics: the number of infected individuals (asymptomatic and symptomatic) at the maximum (when the outbreak is at its most severe), the rate at which the peak infected population is reached (t_max_^-1^), the total infected population after 30 days, and the basic reproductive ratio (*R*_*0*_).

### Basic reproductive ratio

Equations 2.1 - 2.3 give the analytic expression of the basic reproductive ratio (*R*_*0*_) for the model used in this study. This expression for *R*_*0*_ can be deconstructed into two components. Equation 2.2 only contains parameters associated with person to person transmission (*R*_*p*_), while equation 2.3 solely contains parameters associated with transmission from the environment (*R*_*e*_).

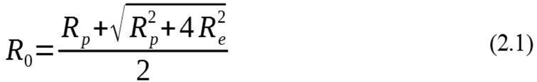

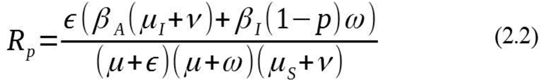

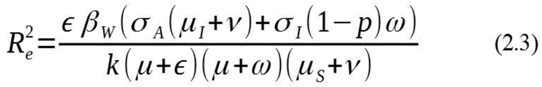

Applying the parameters values in Table 2, the numerical value of the basic reproductive ratio is given as: *R*_*0*_ ∼ 2.82.

## RESULTS

### Model sensitivity analysis

Figure 2 depicts a tornado plot that communicates the sensitivity of the model to permutations in parameters. The model is most sensitive to parameters that are considered virulence-associated (Table 3), and is relatively less sensitive to survival-associated parameters (Table 4).

**Figure 2.**
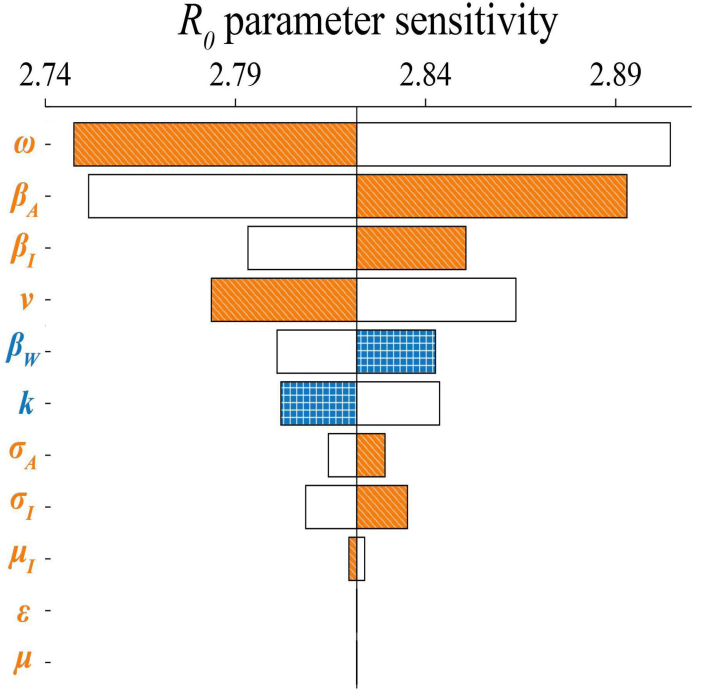
Tornado plot showing the sensitivity of *R*_*0*_ to individual parameter changes. Filled bars indicate the value of *R*_*0*_ when the associated parameter is increased by 5.0% from its nominal value. White bars indicate the value of *R*_*0*_ when the associated parameter is decreased by 5.0%. Blue coloring with checkered patterning indicates a parameter associated with survival, and orange coloring with lined patterning indicates a parameter associated with virulence.

The *R*_*0*_ of the model is most sensitive to the parameters ω, *β*_*A*_, and *v*. The sensitivity of *R*_*0*_ to changes in ω reflects the importance of the rate of conversion into the symptomatic state on model dynamics. In addition, the *β*_*A*_ has a very important influence on the model, consistent with other findings for COVID-19 that have emphasized the importance of asymptomatic transmission in disease spread [22-24].

### Illustrative dynamics of model system

Applying the parameter values in Table 2, Figure 3a demonstrates the base dynamics of the model playing out over the first 100 days while Figure 3b shows the dynamics within the environment over the course of 250 days. In these dynamics, the population begins to be fixed for susceptible hosts. The disease dynamics manifest in the shapes of the curves corresponding to the exposed, asymptomatic, and symptomatic individuals. Note the long tail of the curve corresponding to contamination by the environment. That is, the environment remains infectious even after the infected populations have declined in number. The length and shape of this tail that is influenced by the free-living survival of the virus.

**Figure 3.**
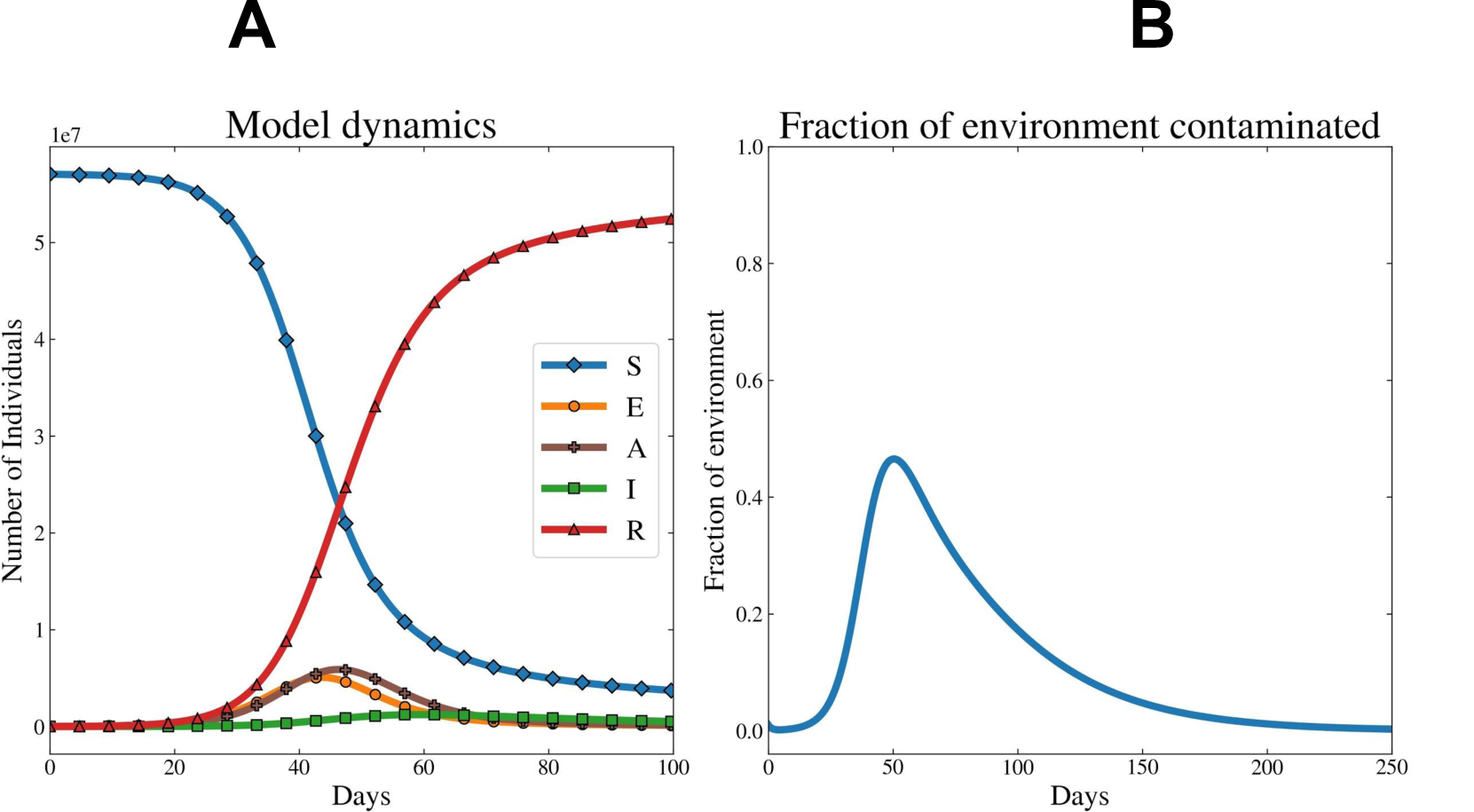
Sample dynamics for the model system. (**A)** Gives the dynamics for all host compartments within the model **(B)** corresponds to the fraction of environmental reservoirs in a setting that are contaminated with infectious virus.

### The epidemic consequences of varying virulence and survival

In the next analysis, we examine the epidemic consequences of varying traits associated with survival and virulence. In Figure 4, we observe how dynamics of the outbreak are influenced by different combinations of traits altering virulence (see Table 3 for a list of virulence-associated parameters) and survival (see Table 4 for a list of survival-associated parameters), changed by +/- 5% (10% overall). In Figure 4D, we demonstrate how changes in virulence and free-living survival traits influence *R*_*0*_, with variation in virulence-related traits having the largest effect on *R*_*0*_. Of note is how the range in *R*_*0*_ values varies widely across virulence-survival values, from nearly 2.0 to 3.7 (Figure 4D).

**Figure 4.**
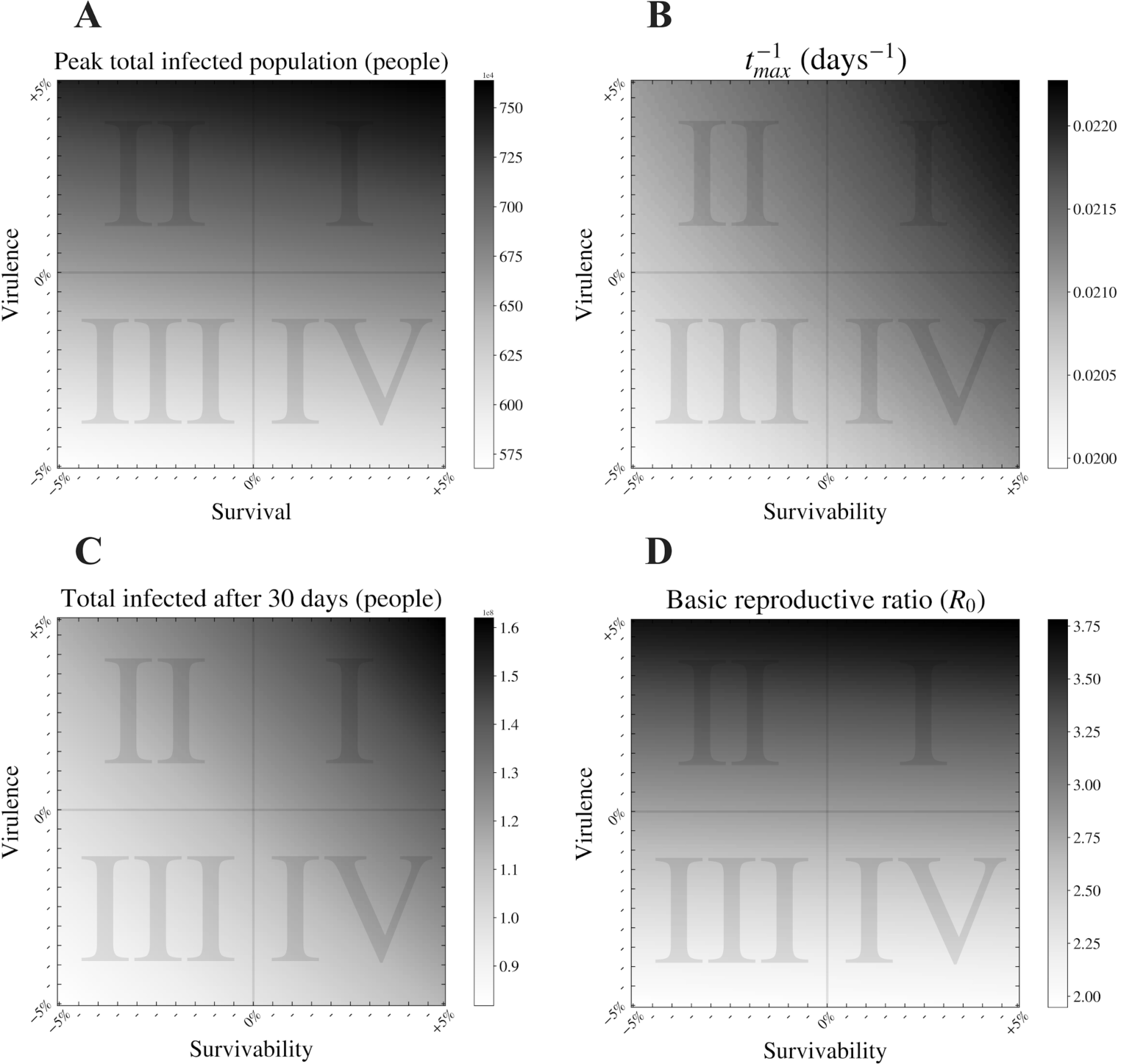
The impact of varying virus virulence and survival assessed on four key epidemic metrics. Here we present heatmaps expressing the change in (A) the number of infected individuals (asymptomatic and symptomatic) at the epidemic peak, (B) the rate at which the epidemic peak is reached, t_max_^-1^, (C) the total infected population after 30 days, and (D) the basic reproductive ratio (*R*_*0*_) when virulence and survival are modulated by +/- 5% within the model.

### Implications of virulence-survival relationships at their relative extremes

Having observed how outbreak dynamics are influenced by variation in traits that alter virulence- survival phenotypes, we then examined how each outbreak metric is influenced by the extreme (+/- 5%) values of the trait combinations considered. Specifically, we assess how a change in pathogen survival affects outbreak dynamics, based on two expected relationships between survival and virulence traits:

- In a positive correlation scenario, high values for survival would be associated with high values for virulence [4,13]. Because the correlations we observe are often not exactly linear, we utilize quadrants to express a trend, allowing for some variance around the expected “line”. In Figure 5, the *positive correlation* scenario can be represented by combinations of virulence and survival residing in quadrants I and III.
- In a negative correlation scenario, high values for survival would be associated with low values for virulence, and low peak total infected population [2,9,15]. Pathogens with a life history that exhibit negative virulence-survival associations that would likely appear in quadrants II and IV in Figure 5.

**Figure 5.**
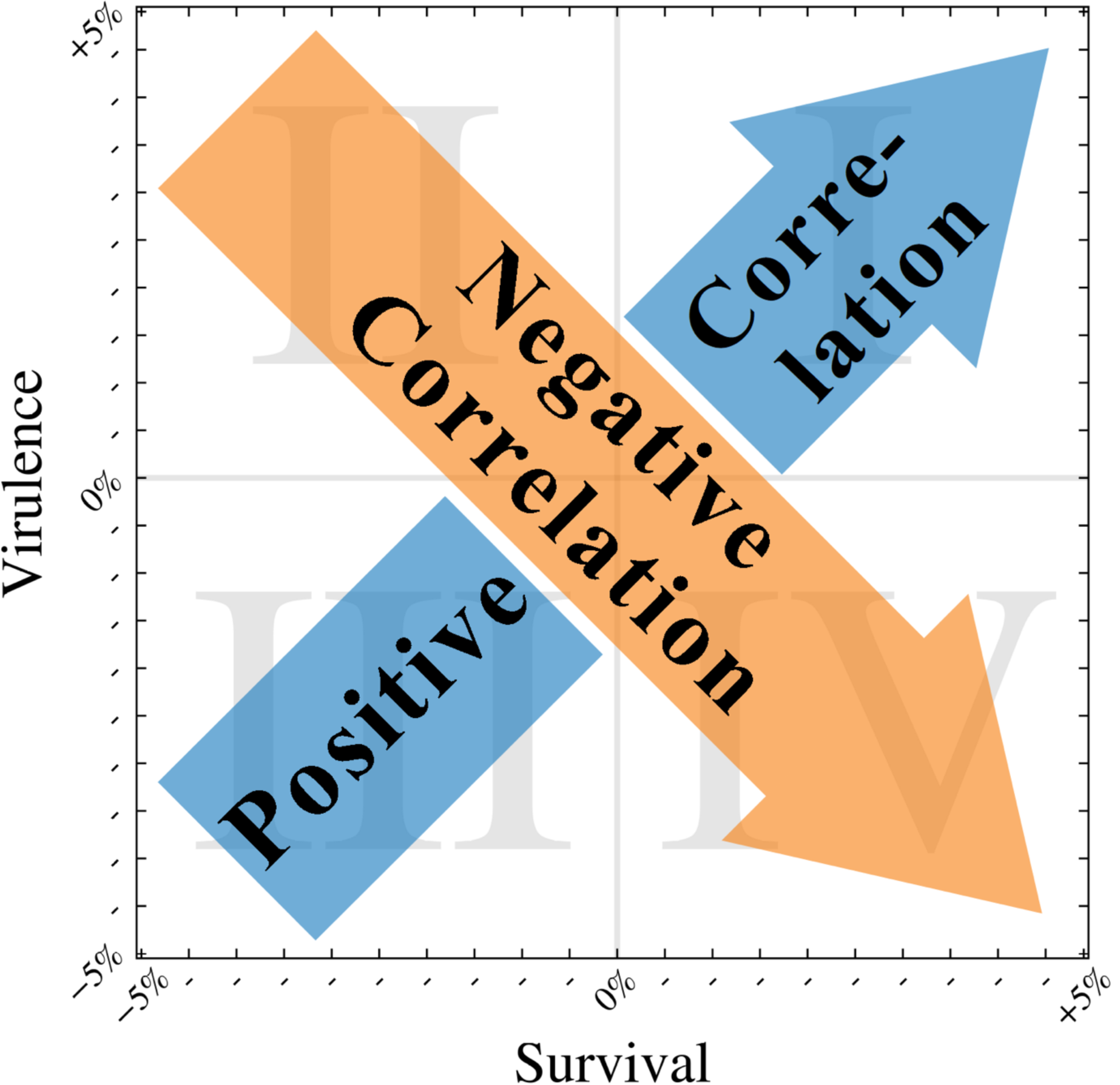
The expected effect of increasing survival on virulence for the two correlation models considered. Here we present a schematic of how the different hypotheses for the relationship between virulence and survival manifest on a map with a structure similar to the heat maps shown in Figure 4. The directions of the arrows depict how increasing survival would affect virulence under the two hypotheses: the blue arrow indicates the flow of an increasing positive correlation dynamic, while the direction of the orange arrow indicates an increasing negative correlation dynamic.

If host-pathogen evolution proceeds according to a positive correlation scenario, all outbreak metrics would show an increase in severity as both survival and virulence increase. Across the range of variation in virulence and survival traits considered (5% above and below the nominal value), peak number of infected individuals increases by ∼35%, the rate at which the peak is reached increases by ∼16%, the total number of infected individuals after 30 days increases by ∼98%, and the *R*_*0*_ increases by ∼94% (Figure 6 and Table 5).

**Table 5.** Comparing epidemic metrics under low survival/low virulence versus high survival/high virulence scenarios (as in the positive correlation *scenario*). For each metric analyzed, these are the heatmap values for the bottom left [at “coordinates” (Vir, Sur) → (−5%, - 5%] and top right [at “coordinates” (Vir, Sur) → (+5%, +5%)] corners.

| **Epidemic**<br>**Metric** | **min virulence,**<br>**min survival** | **max virulence,**<br>**max survival** | **% difference between**<br>**min survival and max**<br>**survival** |
| --- | --- | --- | --- |
| **Peak total**<br>**infected**<br>**(people)** | $5.68 \times 10^6$ | $7.64 \times 10^6$ | +34.51% |
| **$t_{\max}^{-1}$**<br>**(days<sup>-1</sup>)** | $1.99 \times 10^{-02}$ | $2.23 \times 10^{-02}$ | +12.06% |
| **Total after 30**<br>**days (people)** | $8.18 \times 10^{07}$ | $1.62 \times 10^{08}$ | +98.04% |
| **Basic**<br>**reproductive**<br>**ratio ($R_0$)** | 1.95 | 3.78 | +93.84% |

**Figure 6.**
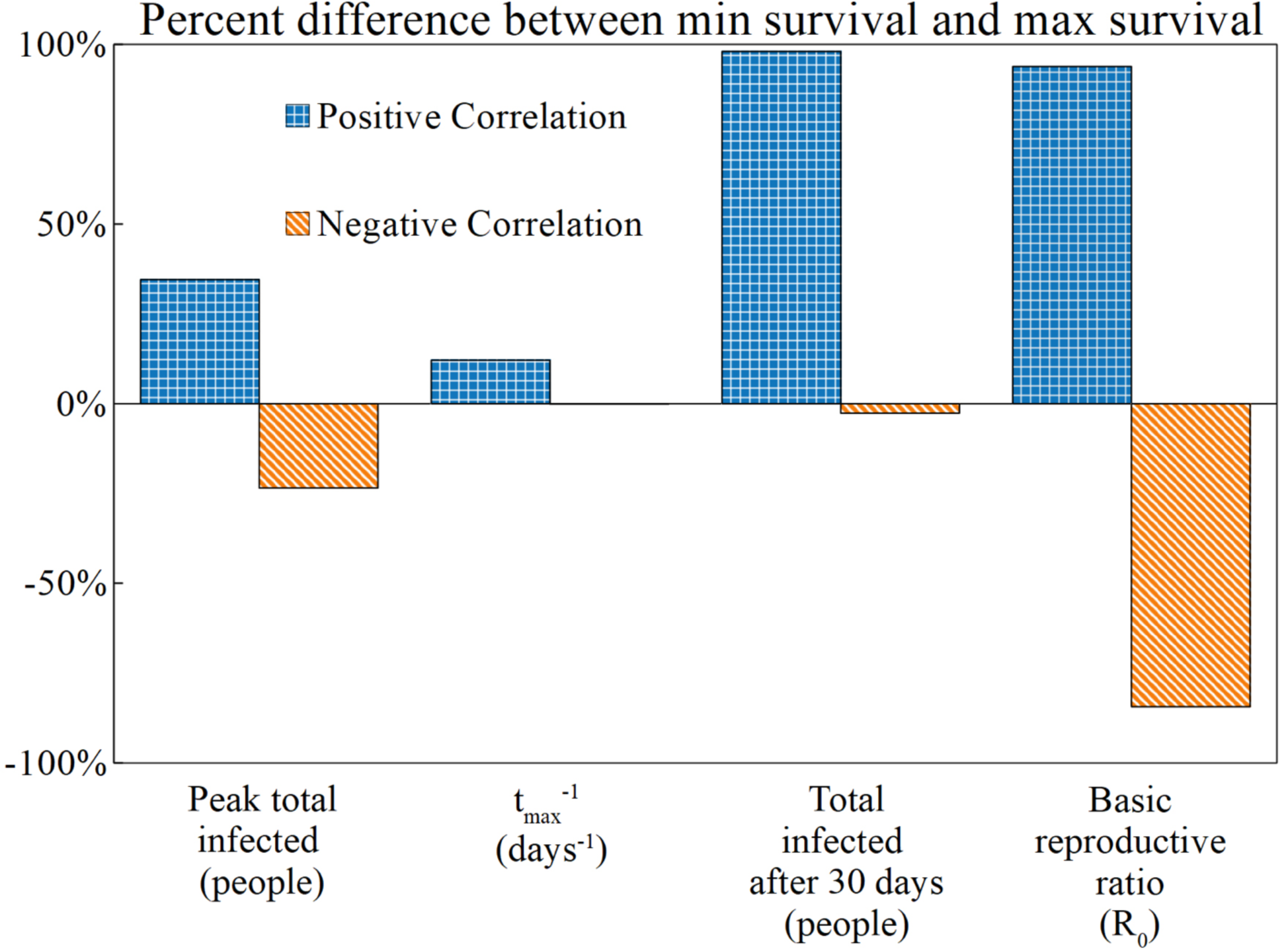
The percent change in SEAIR-W outbreak metrics as survival increases from - 5% to +5% for different virulence-survival relationships (positive correlation and negative correlation). For each metric analyzed we present the percent difference between the minimum and maximum survival values given the two hypotheses tested: (i) positive correlation between survival and virulence) comparing low virulence/low survival to high virulence/high survival) and (ii) negative correlation (high virulence/low survival to high survival/low virulence). The bars here correspond to the values (percent) in the third columns of Table 5 and Table 6, which denote the differences between the minimum and maximum values.

Under negative correlation, outbreak severity generally decreases as survival increases. Across the measured range of variation of virulence-survival traits, the peak number of infected individuals decreases by ∼23%, the rate at which the epidemic peak is reached decreases by 0.15%, the total number of infected individuals decreases by 3%, and the *R*_*0*_ decreases by ∼84% (Figure 6 and Table 6). Across all metrics considered, the effects of increased viral survival on outbreak dynamics is more extreme under the positive correlation than the negative correlation scenario (Figure 6).

**Table 6.** Comparing epidemic metrics under low survival/low virulence versus high survival/high virulence scenarios (as in the negative correlation scenario). For each metric analyzed, these are the global heatmap values for the top left [at “coordinates” (Vir, Sur) → (+5%, −5%)] and bottom right corners [at “coordinates” (Vir, Sur) → (−5%, +5%)]

| Epidemic Metric | max virulence,<br>min survival | min virulence,<br>max survival | % difference between<br>min survival and max<br>survival |
| --- | --- | --- | --- |
| Peak total infected (people) | $7.39 \times 10^6$ | $5.99 \times 10^6$ | -23.47% |
| $t_{\max}^{-1}$ (days<sup>-1</sup>) | $2.10 \times 10^{-02}$ | $2.23 \times 10^{-02}$ | -0.15% |
| Total infected after 30 days (people) | $1.16 \times 10^8$ | $1.13 \times 10^8$ | -2.68% |
| Basic reproductive ratio ($R_0$) | 3.67 | 1.99 | -84.39% |

In Figure 7, we observe the disease dynamics at the survival extremes, and dynamics corresponding to the fraction of the environment that is contaminated with infectious virus. Consistent with the data represented in Figure 6, we observe that minimum and maximum simulations differ more substantially for extreme survival scenarios in the positive correlation scenario than for the negative correlation scenario.

**Figure 7.**
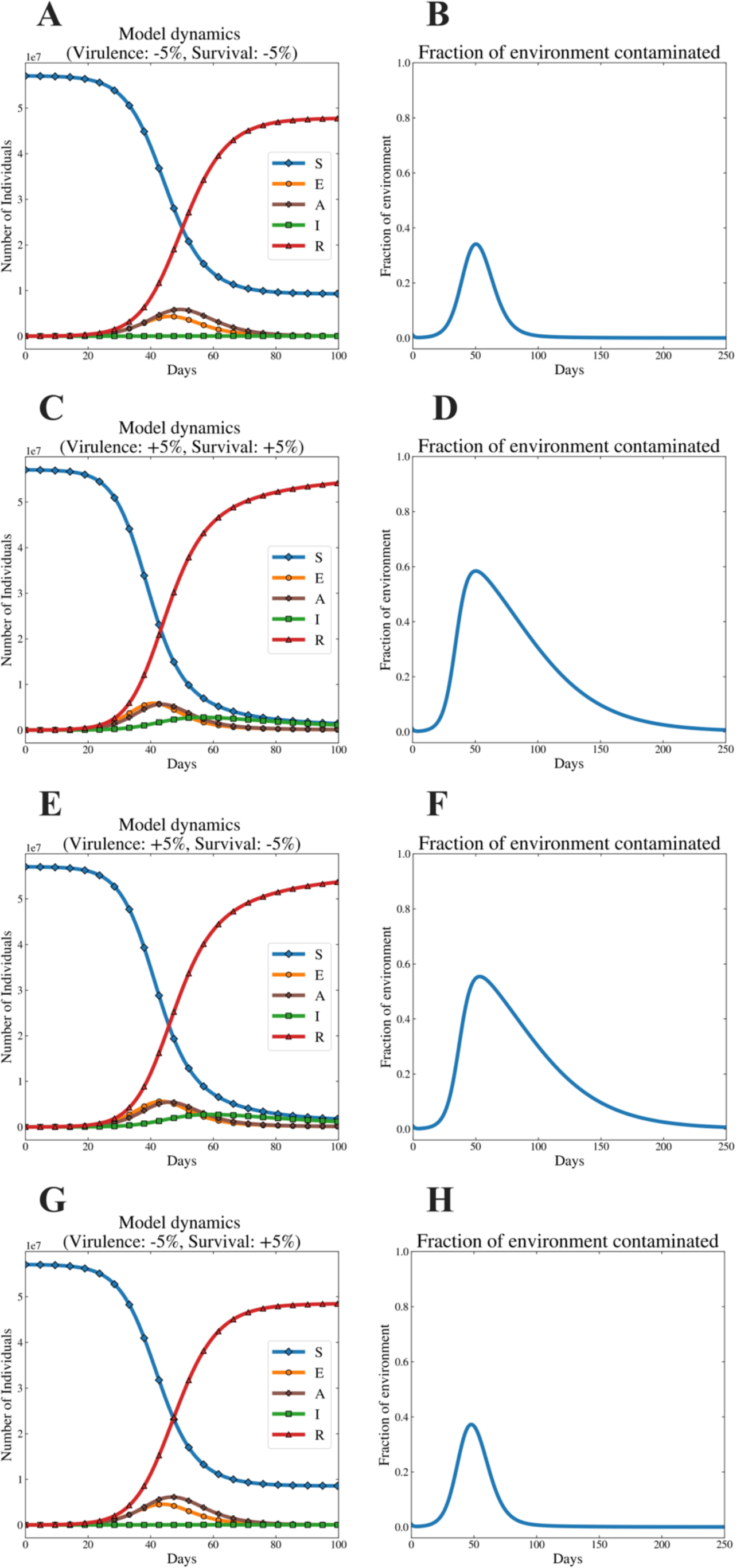
Virus outbreak dynamics for the extreme values of virulence and free-living survival considered for the two different relationships (positive or negative) between virulence and survival traits. These plots are similar to the illustrative dynamics in Figure 1. Here we observe the dynamics of disease corresponding to the extreme values presented in Tables 5 and 6. Figures 7A, C, E, and G depict disease dynamics, and 7B, D, F and H the dynamics of contaminated environments. Figures 7A - D correspond to the parameter values considered for the positive correlation scenario, while Figures 7E - H correspond to the negative correlation scenario.

The feature of the different outbreaks that differs most ostensibly between the correlation scenarios is the time needed to reach the peak number of infected individuals. In the positive correlation simulations, one can observe the low virulence, low survival scenario (Figure 5A and 5B) take longer to reach the peak number of infected individuals. Most notably, however, the low virulence, low survival setting has a far smaller peak of environmental contamination, and shorter tail relative to its high virulence, high survival counterpart (Figure 5D). Similarly, intriguing findings exist in the comparison between the simulation sets corresponding to extremes in the negative correlation setting (Figure 5E-H). Especially notable is the difference in the length of the tail of the environment contamination for the high virulence, low survival combination (Figure 5F) vs. the low virulence, low survival combination variant (Figure 5H). The explanation is that, in this model, higher virulence influences (among many other things) the rate at which virus is shed into the environment from either the asymptomatic (*σ*_*A*_) or symptomatic (*σ*_*I*_) host. We observe how the high virulence, low survival simulation (Figure 5E) features a symptomatic peak that is larger in size, and prolonged relative to the lower virulence counterpart (Figure 5G). This relatively large symptomatic population is shedding infectious virus into the environment for a longer period of time, contributing to the long tail of contaminated environments observed in Figure 5F.

## DISCUSSION

The virulence-survival relationship drives the consequences of virus evolution during an outbreak. In this study, we examined how different virulence-survival relationships may dictate different features of outbreaks as pathogens evolve (according to the positive or negative correlation scenarios). When the parameter space for virulence and survival is mapped, we find that certain outbreak metrics are more sensitive to change in free-living survival traits than others, and that the nature of this sensitivity differs depending on whether survival and virulence are positively or negatively correlated.

For the positive correlation scenario, when free-living survival varies between 5% below and above the nominal value, we observed a dramatic change in the total number of infected individuals in the first 30 days (98% increase from minimum survival to maximum survival), and the *R*_*0*_ nearly doubles (94% increase). These two traits are, of course, connected: the theoretical construction of the *R*_*0*_ metric specifically applies to settings where a pathogen is spreading in a population of susceptible hosts [31, 32].

When survival and virulence are negatively correlated, different outbreak dynamics arise: while the *R*_*0*_ difference between minimum and maximum survival is significant (∼84% decrease), the total number of infected individuals only changes by roughly 3%.

Summarizing, at the extreme survival values in the negative correlation setting, the hypothetical virus population has a reduction in virulence (using either of the stated definitions for virulence). This decreased virulence has a dramatic impact on the *R*_*0*_. Notably, this large difference between the *R*_*0*_ at higher and lower survival values does not translate to a difference in the total number of infected individuals in the first 30 days of an infection (the early outbreak window). That is, in a negative correlation scenario, a highly virulent and less virulent virus population can have similar signatures on a population with respect to the number of infected individuals in the first month. Thus, simply measuring the number of infected individuals in the first month of an outbreak is unlikely to reveal whether not a pathogen population has evolved.

Across scenarios, we observe that relatively small changes in virulence and survival can have large impacts on outbreak dynamics. That the entirety of the results in this study took place within a relatively small range of parameter values (+/− 5%) is significant. This suggests that it doesn’t take large phenotypic changes in free-living survival in order to have a notable imprint on outbreak dynamics.

## Practical implications for the understanding of outbreaks caused by emerging viruses

In summary, our results suggest that carefully constructed, mechanistically sound models of epidemics are important, both for capturing the dynamics of an outbreak, and for understanding how evolution of survival and virulence influence disease dynamics. For example, our results highlight that questions surrounding the evolution of viruses like SARS-CoV-2 during an epidemic require very specific types of evidence. Detecting the consequences of virus evolution would depend on which feature of an outbreak an epidemiologist is measuring: from our analysis, the *R*_*0*_ is most significantly impacted by changes in virulence and survival. In addition, the total number of infected individuals in the early window, and the size of the infected “peak” would each be impacted most readily by changes in virulence-survival traits. The rate at which the epidemic peak was reached, on the other hand, showed relatively little change as survival increased, or between the two correlation scenarios.

The results have practical considerations for how we interpret the potential for local adaptation in pathogens. For example, in the context of COVID-19, speculation has surfaced about the potential mutations in certain virus populations that are associated with increased transmission. As of July 1, 2020, evidence for these types of adaptations is tenuous, and any grand conclusions or predictions are premature. We propose that the process of predicting the evolutionary consequences of viral adaptation should encompass several discrete steps. Firstly, we should determine whether molecular signatures exist for adaptive evolution. Adaptive evolution would manifest in observable differences in genotype and phenotype, perhaps manifesting in the natural history of disease. Secondly, we should aim to attain knowledge of the underlying mechanistic relationship between survival and virulence. This knowledge is not necessarily easy to attain (it requires extensive laboratory studies) but would allow us predictive power: we may be able to extrapolate how changes in some traits influence others. Finally, with this information, mechanistically-informed models can be constructed that may give useful insight into the future trajectory of an outbreak.

While the stochastic, sometimes entropic nature of epidemics renders them very challenging to predict [33], we suggest that canons such as life history theory and the evolution of virulence provide useful lenses that can aid in our ability to interpret how life history changes at the virus level will manifest at the epidemiological scale. More generally, we propose that in an age of great accumulation of genomic and phenotypic data in many pathogen-host systems, we continue to responsibly apply or modify existing theory, in order to collate said data into an organized picture—and potentially paradigm—for how different components of the host-parasite interaction are related to one another, and influence the shape of viral outbreaks of various kinds.

## ACKNOWLEDGEMENTS

The authors would like to thank members of the Ogbunu Lab for helpful discussions and input on all aspects of the manuscript. In particular, we thank MMD and ALM for especially useful interactions.

