## Supplemental Information for "The epidemiological signature of pathogen populations that vary in the relationship between free-living survival and virulence"

**This document contains:**

Supplemental Tables S1-S4

Supplemental Figures S1-S6

Supplemental References

| Symbols | Definitions | Supplemental References |
|---|---|---|
| $\mu$ | Natural Death Rate (Reciprocal of the average life expectancy of 17 countries sampled) | 1, 2 |
| $\mu_S$ | Infected death rate | 3,4 |
| $\eta$ | Incubation Period | 5 |
| $1/\omega$ | Expected time in the asymptomatic state | Fitted and dependent on $\eta$ |
| $\nu$ | Recovery rate (Average of 3 to 6 weeks) | 5 |
| $p$ | Fraction that move along the "mild" recovery track | 6 |
| $k$ | Viral decay rate in environment (using average of all material values, wood, steal, cardboard, plastic) | 7 |

**Table S1. Fixed parameters, and their references.** The mathematical model utilized in this study is derived from an existing one (Ogbunugafor et al., 2020; see main text reference 26). While all references and estimation methods can be found there, here we provide several of the fixed parameters and their references.

*Alternative definition of virulence*. We ran the analysis observed in the main text for a scenario where virulence is only composed of those parameters that directly affect host well-being or fitness. While $\sigma_A$ and $\sigma_s$ are traits included in the main text definition reflect traits that don't directly affect host fitness. Instead, they correspond to traits (the shedding of virus) the consequences of infection. In order to examine the results of our analysis using a stricter definition of virulence, we eliminated these from this alternative analysis. Below find the find the virulence definition and a series of analyses that utilize this alternative definition. The differences between the two definitions can be observed in the side by side comparisons (S4-S6). The conclusions are qualitatively the same

| **Symbols** | **Definition** |
|---|---|
| $\mu_I$ | Infected death rate (Natural death rate + disease induced death rate) |
| $\eta = (\omega - \varepsilon^{-1})$ | SARS-CoV-2 incubation period |
| $1/\omega$ | Rate of transfer from asymptomatic to symptomatic |
| $v$ | Recovery rate (Average of 3 to 6 weeks) |
| p | Percent that move along the "mild" recovery track |
| $\beta_A$ | (Contact rate of people with people) x (transmission probability of people to people by A-person) |
| $\beta_I$ | (Contact rate of people with people) x (transmission probability of people to people by I-person) |
| $1/\varepsilon$ | Average number of days before infectious |

**Table S1. Supplemental virulence definition**. containing only items that directly affect host well-being (fitness). We ran calculations observed in Figure S1-S3 with this different definition.

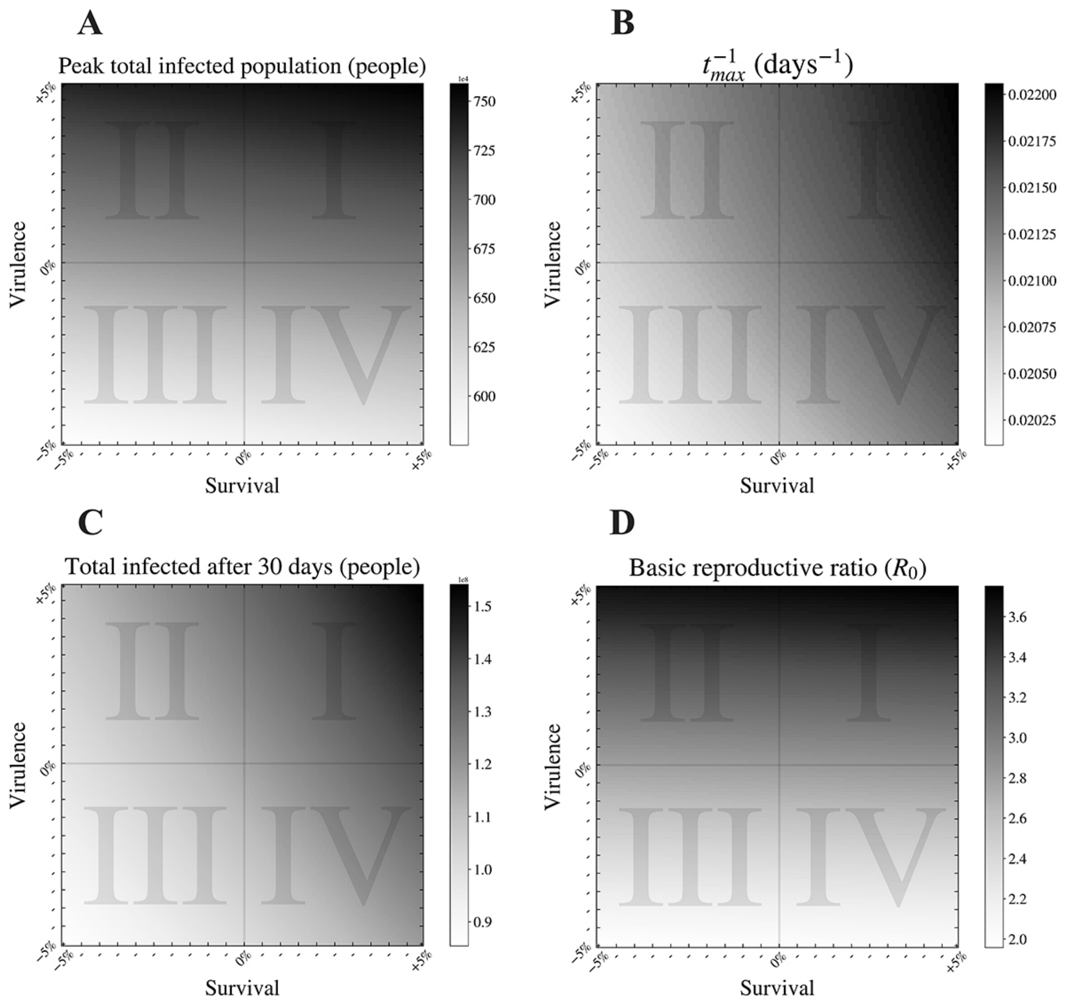


**Figure S1**. **Alternative virulence definition**: **The impact of varying virus virulence and survival assessed on four key epidemic metrics.** As in the main text Figure 4, here we present heatmaps expressing the change in (A) the number of infected individuals (asymptomatic and symptomatic) at the epidemic peak, (B) the rate at which the epidemic peak is reached, $t_{max}^{-1}$, (C) the total infected population after 30 days, and (D) the basic reproductive ratio ($R_0$) when virulence and survival are modulated by +/- 5% within the

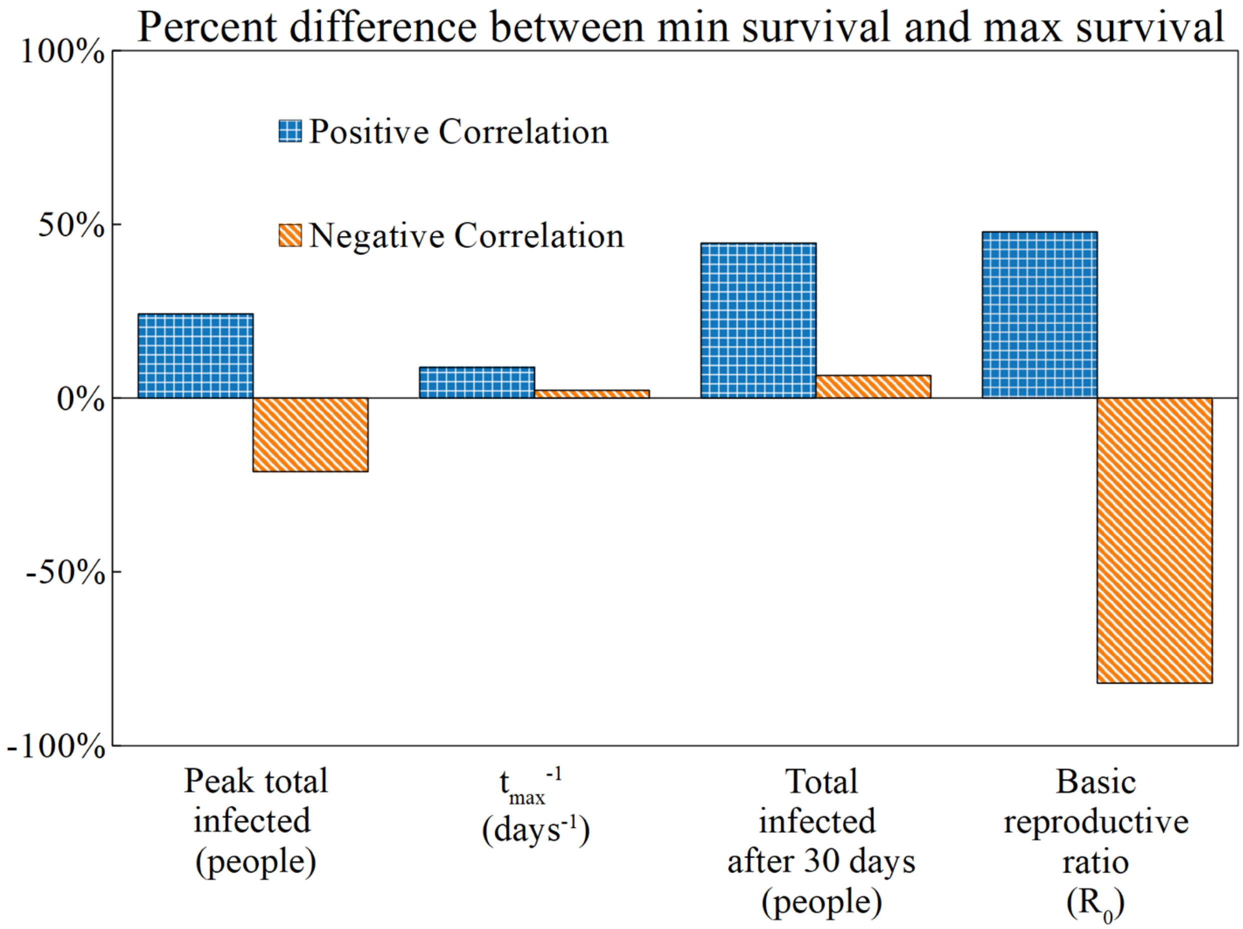


**Figure S2. Alternative virulence definition: The percent change in SEAIR-W outbreak metrics as survival increases from -5% to +5% for different virulence-survival relationships (positive correlation and negative correlation).** As in the main text Figure 6, for each metric analyzed we present the percent difference between the minimum and maximum survival values given the two hypotheses tested: (i) positive correlation between survival and virulence) comparing low virulence/low survival to high virulence/high survival) and (ii) negative correlation (high virulence/low survival to high survival/low virulence). The bars here correspond to the values (percent) in the third columns of Table 5 and Table 6, which denote the differences between the minimum and maximum values.

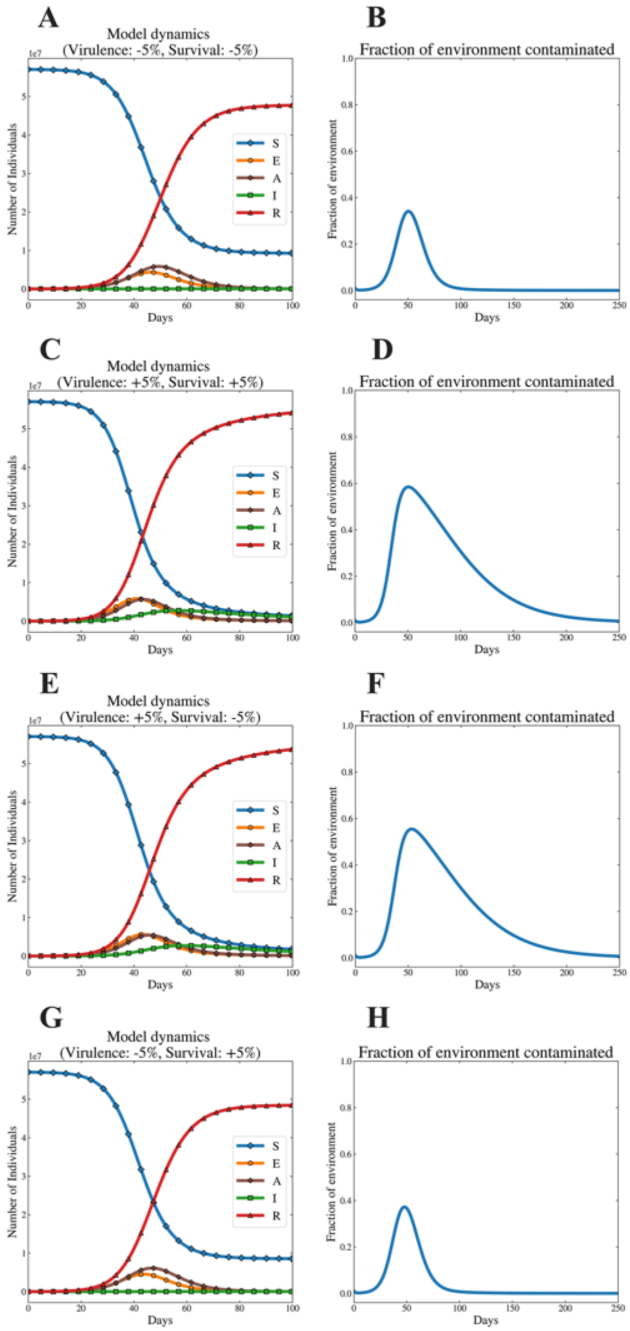


**Figure S3. Alternative virulence definition: Virus outbreak dynamics for the extreme values of virulence and free-living survival considered for the two hypotheses.** As in the main text Figure 7, here we observe the dynamics of disease corresponding to the extreme values presented in Tables 5 and 6. Figures 7A, C, E, and G depict disease dynamics, and 7B, D, F and H the dynamics of contaminated environments. Figures 7A - D correspond to the parameter values considered for the *Curse of the Pharaoh*, while Figures 7E - H correspond to the *Goldilocks hypothesis.*

**positive correlation**

| Epidemic Metric | min virulence, min survival | max virulence, max survival | % difference between min survival and max survival |
|---|---|---|---|
| Peak total infected (people) | 5.75E+06 | 7.59E+06 | 24.24% |
| Tmax-1 (days-1) | 2.01E-02 | 2.21E-02 | 8.81% |
| Total infected after 30 days (people) | 8.54E+07 | 1.54E+08 | 44.59% |
| Basic reproductive ratio (R0) | 1.96 | 3.75 | 47.83% |

**Table S3. Alternative virulence definition: Comparing epidemic metrics under low survival/low virulence versus high survival/high virulence scenarios. As in the main text Table 5, f**or each metric analyzed, these are the heatmap values for the bottom left [ at "coordinates" (Vir , Sur ) → ( -5% , -5% ) ] and top right [ at "coordinates" ( Vir , Sur ) → ( +5% , +5% ) ] corners.

**negative correlation**

| Epidemic Metric | max virulence, min survival | min virulence, max survival | % difference between min survival and max survival |
|---|---|---|---|
| Peak total infected (people) | 7.35E+06 | 6.07E+06 | -21.13% |
| Tmax-1 (days-1) | 2.09E-02 | 2.13E-02 | 2.28% |
| Total infected after 30 days (people) | 1.11E+08 | 1.18E+08 | 6.53% |
| Basic reproductive ratio (R0) | 3.64 | 2.00 | -82.04% |

**Table S4. Alternative virulence definition: Comparing epidemic metrics under low survival/low virulence versus high survival/high virulence scenarios (as in the *negative correlation scenario*).** For each metric analysed, these are the global heatmap values for the top left [ at "coordinates" (Vir , Sur) → (+5% , -5%) ] and bottom right corners [at "coordinates" ( Vir , Sur ) → (-5% , +5% )]

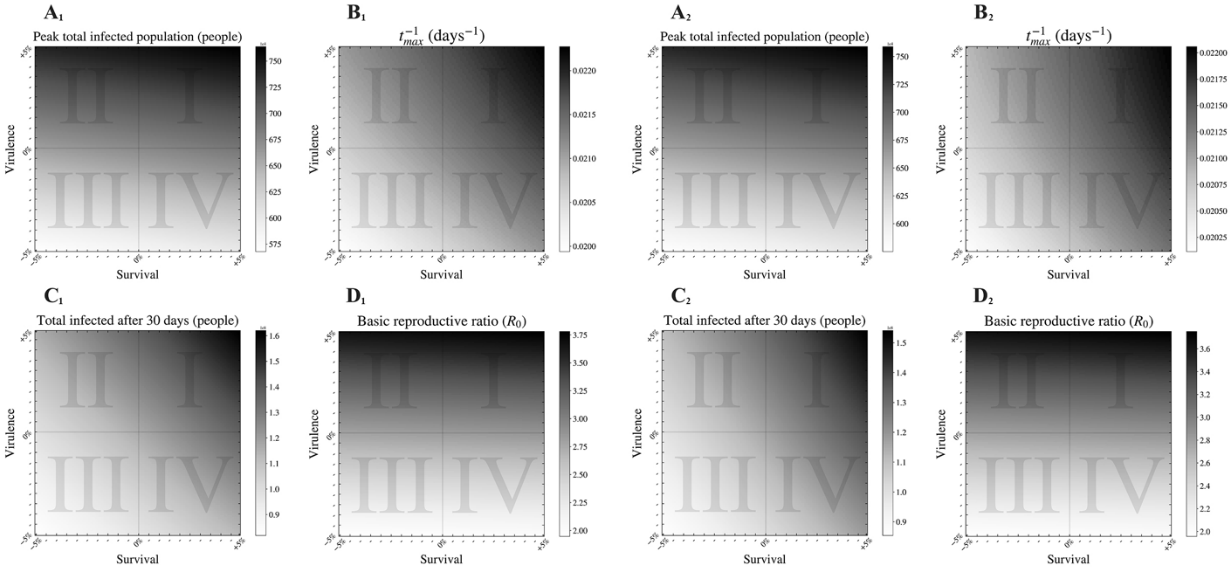


**Figure S4.** Side by side comparison of the main text virulence definition (A1 → D1) and alternative definition of virulence (A2 → D2).

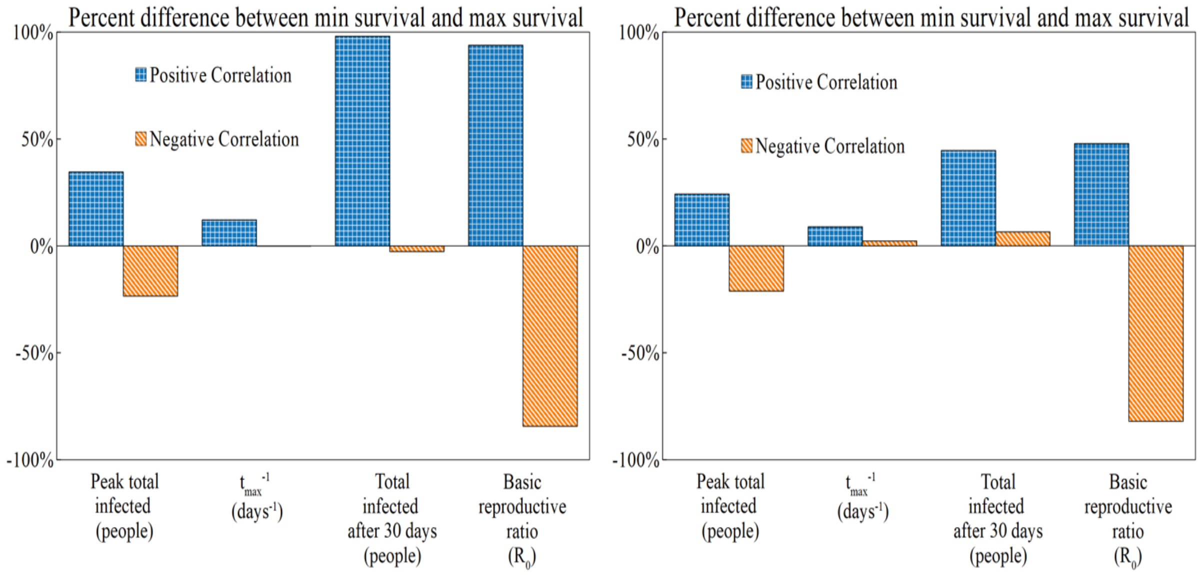


**Figure S5.** Side by side comparison of the main text virulence definition (left) and alternative definition of virulence (right).

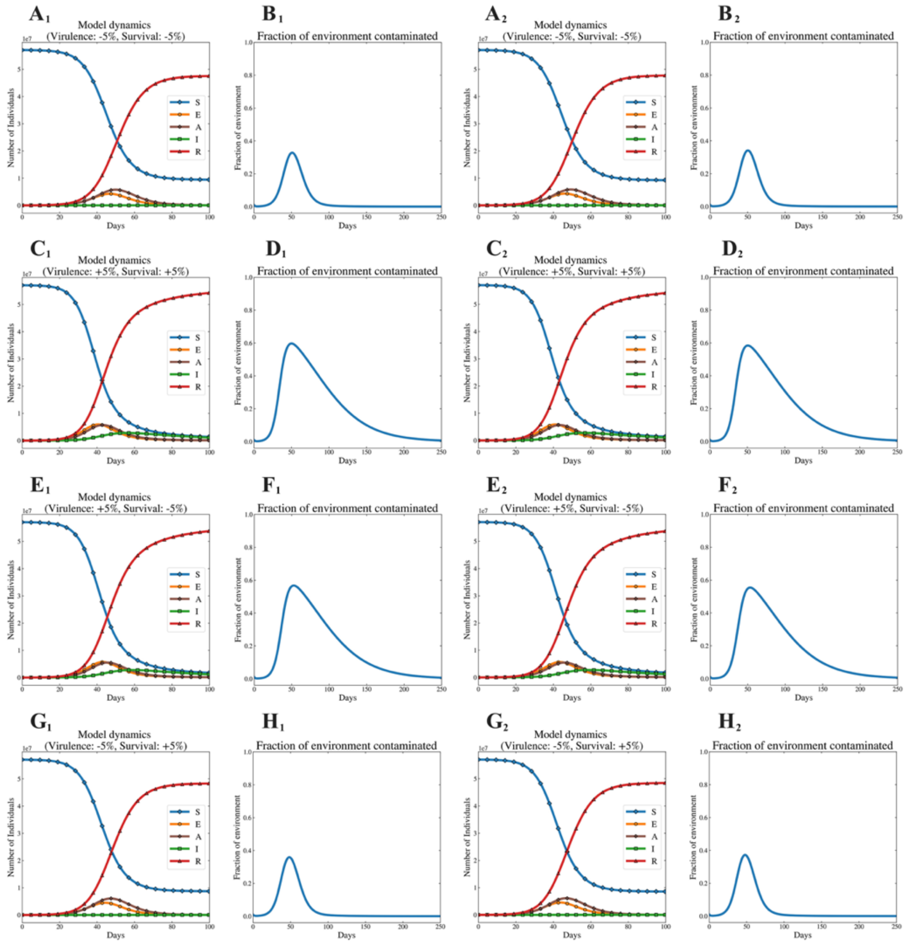


**Figure S6.** Side by side comparison of the main text definition (A1 → H1) and supplemental definition of virulence ($A_2$ → $H_2$).
